# Oncogenic ERRB2 signals through the AP-1 transcription factor to control mesenchymal-like properties of oesophageal adenocarcinoma

**DOI:** 10.1101/2022.08.14.503893

**Authors:** Samuel Ogden, Ibrahim Ahmed, Paul Fullwood, the OCCAMS consortium, Chiara Francavilla, Andrew D. Sharrocks

**Author notes:** Corresponding authors: Chiara Francavilla, Andrew Sharrocks (lead contact).

## Abstract

Oesophageal adenocarcinoma (OAC) is a deadly disease with poor survival statistics and few targeted therapies available. One of the most common molecular aberrations in OAC is amplification or activation of the gene encoding the receptor tyrosine kinase ERBB2, and ERBB2 is targeted in the clinic for this subset of patients. However, the downstream consequences of these ERBB2 activating events are not well understood. Here we used a combination of phosphoproteomics, open chromatin profiling and transcriptome analysis on cell line models and patient-derived datasets to interrogate the molecular pathways operating downstream from ERBB2. Integrated analysis of these data sets converge on a model where dysregulated ERBB2 signalling is mediated at the transcriptional level by the transcription factor AP-1. AP-1 in turn controls cell behaviour by acting on cohorts of genes that regulate cell migration and adhesion, features often associated with EMT. Our study therefore provides a valuable resource for the cancer cell signalling community and reveals novel molecular determinants underlying the dysregulated behaviour of OAC cells.

## Introduction

The RAS-ERK pathway and its upstream regulators is one of the most commonly disrupted pathways in cancer and its hyperactivation through oncogenic mutations drives the tumourigenic phenotype (1, 2). Hyperactive RAS-ERK pathway signalling has major impacts in the nucleus on the activity of gene regulatory networks and the underlying regulatory chromatin landscape. Multiple transcription factors and chromatin remodelling proteins have been shown to be direct ERK targets (reviewed in 3). In common with many other cancers, the RAS-ERK pathway is frequently activated in oesophageal adenocarcinoma (OAC) with multiple components being mutationally activated in different patients (60%-76%)(4,5). One of the most commonly mutated genes encodes the upstream receptor tyrosine kinase ERBB2, whose locus is amplified in up to 32% of cases (4,5). Patients harbouring these mutations can be treated with the ERBB2 targeting antibody trastuzumab, one of the few targeted therapies available for OAC patients (6; reviewed in 7). Indeed, due to usual late diagnosis and the lack of subsequent targeted therapies available, the current prognosis for OAC patients is poor with 5 year survival rates being only ~20% (8). However, our understanding of the molecular and cellular consequences of ERBB2 activation in OAC are poorly understood. Similarly, we know little about the molecular outcomes caused by the activation of alternative RTKs and commonly used downstream pathway components that are mutated in other OAC patients. To bridge this knowledge gap, we recently studied the effects of pharmacological ERBB2 inhibition on the gene regulatory pathways operational in OAC cells as they transition to a drug resistant state (9). In that study we focussed on the programmes that become newly operational following ERBB2 inhibition, and demonstrated a role for the transcription factor HNF4A and also the transcriptional co-activator PPARGC1A in rewiring the metabolic pathways to allow cells to survive and subsequently acquire a resistant state.

In this study, we extended our investigation of ERBB2 function and focussed on the immediate consequences of inhibitor treatment to identify the pathways and processes controlled by ERBB2 in OAC cells. By combining phosphoproteomics, chromatin accessibility mapping and transcriptomics, we uncovered the transcription factor AP-1 as a major mediator of ERBB2 activity in OAC. This is consistent with the finding that some of the major transcription factor targets of RTK signalling are in the AP-1 family. Individual subunits can be phosphorylated and activated (10, 11, 12) and/or their levels increased at the transcriptional level following pathway activation (reviewed in 13). AP-1 itself controls multiple cellular processes (14), with its connections to *CCND1* expression (15) linking it to cell cycle control and cell proliferation, one of the key cancer cell attributes (16). However, rather than an expected role in cell proliferation, AP-1 instead mediates ERBB2 signalling effects in OAC through controlling EMT-related properties such as actin cytoskeleton re-organisation and cell migration.

## Materials and methods

### Cell culture and treatments

OE19, and ESO26 cells were cultured in RPMI 1640 (ThermoFisher Scientific, 52400) supplemented with 10% foetal bovine serum (ThermoFisher Scientific, 10270) and 1% penicillin/streptomycin (ThermoFisher Scientific, 15140122). KYAE1 cells were cultured in 1:1 RPMI 1640:F12 (ThermoFisher, 11765054) supplemented with 10% foetal bovine serum and 1% penicillin/streptomycin. HEK293T cells were cultured in DMEM (ThermoFisher Scientific, 22320-022) supplemented with 10% foetal bovine serum. Cell lines were cultured at 37°C, 5% CO_2_ in a humidified incubator.

### Cell growth assays

Crystal violet assays were performed as described previously (9).

### Dominant negative FOS over-expression

pINDUCER20-GFP-AFOS (ADS5006) (17) was packaged into lentivirus and OE19, KYAE1 and ESO26 cells were then transduced with lentivirus as described previously (9). Polyclonal cells were selected for 2 weeks using 250 μg/mL G418 (ThermoFisher Scientific, 10131027). Transduced cells were then seeded at 2×10^4^ cells / cm^2^ for subsequent experiments. 24 hours later, dominant negative FOS (acidic FOS, aFOS; 18) was induced by doxycycline treatment.

### SILAC labelling

OE19 cells were labelled in SILAC RPMI (Life Technologies, 88365) supplemented with 10% dialysed fetal bovine serum (Sigma, F0392) and 1% penicillin/streptomycin for 20 days (at least 10 cell doublings) to ensure complete incorporation of amino acids. Media was supplemented with labelled amino acids: light condition [Lys0 (Sigma, L8662), Arg0 (Sigma, A6969)], medium condition [Lys4 (CIL, DLM-2640-PK), Arg6 (CIL, CLM-2265-H-PK), heavy condition [Lys8 (CIL, CNLM-291-H-PK), Arg10 (CIL, CNLM-539-H-PK)].

### Mass spectrometry-based proteomics and phosphoproteomics

Sample preparation: SILAC-labelled OE19 cells were seeded at 1.92×10^4^ cells/cm^2^. In total 2.4×10^7^ cells were seeded for light and medium conditions, and 3.16×10^7^ cells were seeded for the heavy condition to account for cell death caused by lapatinib treatment. 24 hours after seeding, the light condition was treated with DMSO for 2 hours, medium condition was treated with 500 nM lapatinib (Selleckchem, S1028) for 2 hours and the heavy condition was treated with 500 nM lapatinib for 24 hours. Cells were then washed with PBS and lysed at 4°C in ice-cold 1% triton lysis buffer supplemented with protease inhibitors (ThermoFisher Scientific, A32963) and phosphatase inhibitors (5 nM Na_3_VO_4_, 5 nM NaF, 5 mM β-glycerophosphate). Samples were prepared as described (Smith et al., 2021). Briefly, a four-fold excess of ice-cold acetone was added and proteins were precipitated overnight at −20°C. Precipitated protein was solubilised in denaturation buffer (10 mM HEPES, pH 8.0, 6 M urea, 2 M thiourea) and proteins were quantified using a Bradford assay (Pierce, 23200). Three mg of each light, medium and heavy replicate were combined, resulting in 9 mg of total proteins. Cysteines were reduced using 1 mM dithiothreitol (DTT) and alkylated with 5.5 M chloroacetamide (CAA). For proteome analysis alkylated proteins were resolved by SDS-PAGE (8-12%, Invitrogen) and then fixed in gel and visualized with Colloidal Blue staining (Invitrogen, LC6025). Gel lanes were separated into eight slices, minced and destained with 50% ethanol in acetonitrile. Proteins were digested with sequencing grade, modified trypsin (Sigma, TRYPSEQM-RO) followed by quenching with 1% trifluoroacetic acid. For the phosphoproteome, alkylated proteins were digested using Lysyl Endopeptidase^®^ (FUJIFILM Wako Pure Chemical Corporation, 125-05061) and sequencing grade modified trypsin (Sigma, TRYPSEQM-RO) followed by quenching with 1% trifluoroacetic acid. Peptides were purified using reversed-phase Sep-Pak C18 cartridges (Waters, USA) and eluted with 50% acetonitrile. 6 mL of 12% trifluoroacetic acid was added to the eluted peptides and subsequently enriched with TiO_2_ beads (5 μm, GL Sciences Inc., Tokyo, Japan). The beads were suspended in 20 mg / mL 2,5-dihydroxybenzoic acid, 80% acetonitrile, 5% trifluoroacetic acid and the samples were incubated in a sample to bead ratio of 1:2 (w/w) in batch mode for 15 minutes with rotation. After 5 minutes centrifugation the supernatant was collected and incubated a second time with a two-fold dilution of the previous bead suspension. Beads were washed with 10% acetonitrile, 6% trifluoroacetic acid followed by 40% acetonitrile, 6% trifluoroacetic acid and collected on C8 STAGE-tips (Waters, USA) and finally washed by 80% acetonitrile, 6% trifluoroacetic acid. Phosphorylated peptides were eluted using 20 μL 5% NH_3_ followed by 20 μL 10% NH_3_ in 25% acetonitrile. Phosphorylated peptides were evaporated to a final volume of 5 μL in a SpeedVac. Concentrated phosphorylated peptides were acidified with addition of 20 μL 0.1% trifluoroacetic acid, 5% acetonitrile and loaded on C18 STAGE-tips. Peptides were eluted from STAGE-tips with 20 μL of 40% acetonitrile followed by 10 μL 60% acetonitrile. Peptides were then reduced to a final, 5 μL volume by SpeedVac and 5 μL 0.1% formic acid, 5% acetonitrile was added. Samples were analysed by LC-MS/MS using a QE HF (ThermoFisher Scientific).

Mass Spectrometry: Purified peptides were analysed by LC-MS/MS using an UltiMate® 3000 Rapid Separation LC (RSLC, Dionex Corporation, Sunnyvale, CA) coupled to a QE-HF (Thermo Fisher Scientific, Waltham, MA) mass spectrometer (19). Mobile phase A was 0.1% FA in water and mobile phase B was 0.1% FA in ACN and the column was a 75 mm x 250 μm inner diameter 1.7 mM CSH C18, analytical column (Waters). Peptides were separated using a gradient that went from 7% to 18% B in 64 min., then from 18% to 27% B in 8 min. and finally from 27% B to 60% B in 1 min. The column was washed at 60% B for 3 min. and then re-equilibrated for a further 6.5 minutes. At 85 minutes the flow was increased to 300 nl/min until the end of the run at 90min. Mass spectrometry data was acquired in a data directed manner for 90 min. in positive mode. Peptides were selected for fragmentation automatically by data dependent analysis on a basis of the top 8 (phosphoproteome analysis) or top 12 (proteome analysis) with m/z between 300 to 1750Th and a charge state of 2, 3 or 4 with a dynamic exclusion set at 15 sec. The MS Resolution was set at 120,000 with an AGC target of 3e6 and a maximum fill time set at 20ms. The MS2 Resolution was set to 60,000, with an AGC target of 2e5, and a maximum fill time of 110 ms for Top12 methods, and 30,000, with an AGC target of 2e5, and a maximum fill time of 45 ms for Top8 analysis. The isolation window was of 1.3Th and the collision energy was of 28.

Raw Files Analysis: Raw data were analysed by the MaxQuant software suite (https://www.maxquant.org; version 1.6.2.6) using the integrated Andromeda search engine (20). Proteins were identified by searching the HCD-MS/MS peak lists against a target/decoy version of the human Uniprot Knowledgebase database that consisted of the complete proteome sets and isoforms (v.2019; https://uniprot.org/proteomes/UP000005640_9606) supplemented with commonly observed contaminants such as porcine trypsin and bovine serum proteins. Tandem mass spectra were initially matched with a mass tolerance of 7 ppm on precursor masses and 0.02 Da or 20 ppm for fragment ions. Cysteine carbamidomethylation was searched as a fixed modification. Protein N-acetylation, N-pyro-glutamine, oxidized methionine, and phosphorylation of serine, threonine, and tyrosine were searched as variable modifications for the phosphoproteomes. Protein N-acetylation, oxidized methionine and deamidation of asparagine and glutamine were searched as variable modifications for the proteome experiments. False discovery rate was set to 0.01 for peptides, proteins, and modification sites. Minimal peptide length was six amino acids. Site localization probabilities were calculated by MaxQuant using the PTM scoring algorithm (21).The dataset were filtered by posterior error probability to achieve a false discovery rate below 1% for peptides, proteins and modification sites. Only peptides with Andromeda score ><u>40 were included.</u>

#### Data and Statistical Analysis

All statistical and bioinformatics analyses were done using the freely available software Perseus, version 1.6.2.1. (22), R framework and Bioconductor 23). We removed proteins or phosphorylated peptides that were not detected in all conditions or all biological replicates. We considered proteins or phosphorylated peptides up- or down-regulated if all 3 biological replicates were enriched by a 1.2 linear fold change (19). Metascape (24) was used for GO analysis. Kinase-Substrate Enrichment Analysis (25) was used for Kinase activity predictions.

### RT-qPCR

RT-qPCR was performed as described previously (9). Primers used are listed in Supplementary Table S6.

### Western blots

Western blots were performed as described previously (9). Antibodies used: anti-GFP (Santa Cruz, sc-8334, 1:2,000), anti-ERK (Cell Signaling Technologies, 4695S, 1:1,000).

### Cell migration and adhesion assays

Cell adhesion assays were performed as previously described (26). Briefly, OE19/KYAE-1-dnFOS cells were treated with 1 μg/mL doxycycline or DMSO control for 24 hours to induce dnFOS. 200,000 cells were then harvested and re-plated for 12 hours at 37 °C, 5% CO_2_ to allow adhesion. Plates were gently tapped and washed with PBS to remove floating cells. Adherent cells were fixed with 4% paraformaldehyde for 10 minutes and stained using 0.1% crystal violet (Sigma-Aldrich, HT90132) for 30 minutes at room temperature followed by extensive washing with water. Plates were allowed to dry and dye solubilised in 10% acetic acid for 10 minutes at room temperature with gentle shaking. Absorbance readings were taken at 570 nm on a SPECTROstar Nano Micoplate Reader (BMG LABTECH).

Cell migration assays were performed as previously described (27). Briefly, OE19/KYAE-1-dnFOS cells were treated with 1 μg/mL doxycycline or DMSO control for 24 hours to induce dnFOS. 200,000 cells were harvested and plated into 8.0 μm transwell inserts (Corning, 353097) in 250 μL serum-free medium. Wells were filled with 750 μL medium containing 20% FBS, 1 μg/mL doxycycline and 5 ng/uL TGF-β. After 24 hours migration at 37 °C, 5% CO_2_, non-migrating cells were removed by wiping with a cotton swab. Migrated cells were fixed with 4% paraformaldehyde for 10 minutes and stained using 0.1% crystal violet. Transwell inserts were dried and imaged using a light microscope. Images were quantified using ImageJ software.

### RNA-sequencing

dnFOS was induced in OE19-dnFOS (28) and KYAE1-dnFOS cells for 48 hours before RNA was isolated and sequenced and analysed as described previously (9). Differentially expressed genes were determined using DESeq2 (29). Approximate FPKMs for plotting purposes were calculated from normalised FPMs output from DESeq2 by dividing the FPMs by the gene length in kilobases (output from featureCounts; 30). Differentially expressed genes with a mean log_2_(FPKM+1) <1 were excluded from the results.

ERBB2 positive OAC samples (*ERBB2*^HIGH^) were determined based on these samples having expression of *ERBB2* greater than the median *ERBB2* expression +2 SD. To identify genes differentially expressed in ERBB2 positive OAC these samples were compared to non-dysplastic Barrett’s oesophagus samples (31).

Metascape (24) was used for gene ontology analysis of differentially expressed genes. Ingenuity Pathway Analysis (32) was used to predict upstream regulators. Gene set enrichment analysis (GSEA) was performed on differentially expressed genes using GSEA v4.0.3 (33), MSigDB Hallmarks v7.2 (34) with the permutation type set to gene set. OAC partial EMT (pEMT) signature genes were obtained from source data from Tyler and Tirosh, 2021 (35). Only genes more specifically expressed in cancer cells relative to the stromal compartment were used.

### ATAC-seq analysis

Amplifications were called in OAC patients as described previously using a custom R script (36). T_007, TCGA-M9-A5M8 and TCGA-IC-A6RE were called as having *ERBB2* amplifications. Amplifications of *ERBB2* in the two TCGA samples were confirmed using cBioPortal (37,38) and visual inspection of the *ERBB2* locus in a genome browser. Initial ATAC-seq data processing was performed as described previously (9). A union peakset was formed of all Barrett’s oesophagus and OAC patients samples, using HOMER v4.9 mergePeaks.pl -d 250 (39) as described previously (40). Regions amplified in any OAC patient sample were removed using BEDtools v2.27.1 intersectBed (41). Differentially accessible peaks were then identified using featureCounts v1.6.2 and DESeq2 v1.14.1 (FDR < 0.05).

Distal regulatory regions (i.e. non-promoter regions) were defined as described previously (40). HOMER v4.9 38) was used for both de novo and ‘known’ transcription factor motif enrichment analysis. HOMER v4.9 annotatePeaks.pl or the basal extension model (GREAT; 42) were used to annotate peaks to genes. Genome browser data was visualised using IGV v2.7.2 (43). Heatmaps and tag density plots of epigenomic data were generated using deepTools (44). To visualise epigenomic data at peaks containing the AP-1 motif, HOMER v4.9 annotatePeaks.pl -size 200 -m was used.

### Bioinformatics

Morpheus (https://software.broadinstitute.org/morpheus/) was used to generate heatmaps. The prcomp function in R v3.6.0 was used for principal component analysis (PCA). Eulerr or ggvenn packages in R v3.6.0 were used for generating Venn diagrams.

### Datasets

All data was obtained from ArrayExpress, unless stated otherwise. Human tissue RNA-seq data was obtained from: E-MTAB-4054 (30) and the OCCAMS consortium (European Genome-Phenome Archive, EGAD00001007496). Human tissue ATAC-seq data was obtained from: E-MTAB-5169 (17), E-MTAB-6751 (40), E-MTAB-8447 (36) and The Cancer Genome Atlas OAC ATAC-seq data were obtained from the GDC data portal (portal.gdc.cancer.gov; 45).

OE19 HNF4A and H3K27ac ChIP-seq was obtained from: E-MTAB-10319. OE19 lapatinib ATAC and RNA-seq was obtained from E-MTAB-10302, E-MTAB-10304. Lapatinib treatment of WTSI-OESO_009, ESO26, KYAE1 and NCI-N87 cells ATAC-seq was obtained from: E-MTAB-10306, E-MTAB-10307, E-MTAB-310, E-MTAB-313 (9). OE19 siERBB2 ATAC-seq and RNA-seq was obtained from: E-MTAB-8576, E-MTAB-8579. KLF5 ChIP-seq data was obtained from E-MTAB-8568 (36).

### Data access

OE19 (28) and KYAE1 dnFOS RNA-seq have been deposited at ArrayExpress (E-MTAB-10319).

The mass spectrometry proteomics data in Thermo Scientific’s *.raw format have been deposited to the ProteomeXchange Consortium via the PRIDE (46) partner repository with the following dataset identifiers:

#### Project Name

ERBB2 signalling and functions in oesophageal cancer. Project accession: PXD029180. Reviewer account details. Username. Password: TUwedE2w.

## Results

### Phosphoproteomic analysis of ERBB2 signalling in OAC cells

To uncover the signalling pathways and downstream targets controlled by oncogenic ERBB2, we treated OE19 OAC cells (which contain *ERBB2* locus amplification; 47) with the ERBB2 inhibitor lapatinib for 2 hrs (to study immediate effects) and 24 hrs (to study subsequent/longer term effects) and analysed changes in the phosphoproteome by SILAC-based quantitative mass spectrometry (48) (Fig. 1A). Changes in protein abundance and phosphorylation levels were determined by calculating the ratios of medium (2 hrs lapatinib) and heavy (24 hrs lapatinib) cell culture conditions to the light condition (control cells). We detected a total of 3,366 phosphorylated sites and 2,318 proteins, combining to produce 1,595 phosphorylated proteins (Supplementary Fig. S1A; Supplementary Tables S1 and S2). The majority of the detected phosphorylated sites (median Andromeda score: 91.2; Supplementary Fig. S1B) were on serine residues (88%) and only 13% of phosphopeptides contained more than one phosphorylated amino acid (Supplementary Fig. S1C), consistent with previous work (19). The replicates from each of the experiments clustered together although one of the 24 hour lapatinib treated samples showed lower reproducibility in the proteome (Supplementary Fig. S1D). At the total protein level, roughly equal numbers of proteins were up- and down-regulated at each time point of lapatinib treatment (Supplementary Fig. S1E). However, as might be expected, reductions in phosphorylation events were more common than increases following lapatinib treatment, with 367 (after 2hrs) and 467 (after 24 hrs) proteins showing decreased phosphorylation (corresponding to 510 and 689 phosphorylated sites respectively) (Supplementary Fig. S1E). Changes in protein level expression generally correlated well with the changes we observed at the mRNA level in RNAseq analysis after 24 hrs (Supplementary Fig. S1F). Overall, these results reveal extensive rewiring of the phosphoproteome following ERBB2 inhibition.

**Figure 1.**
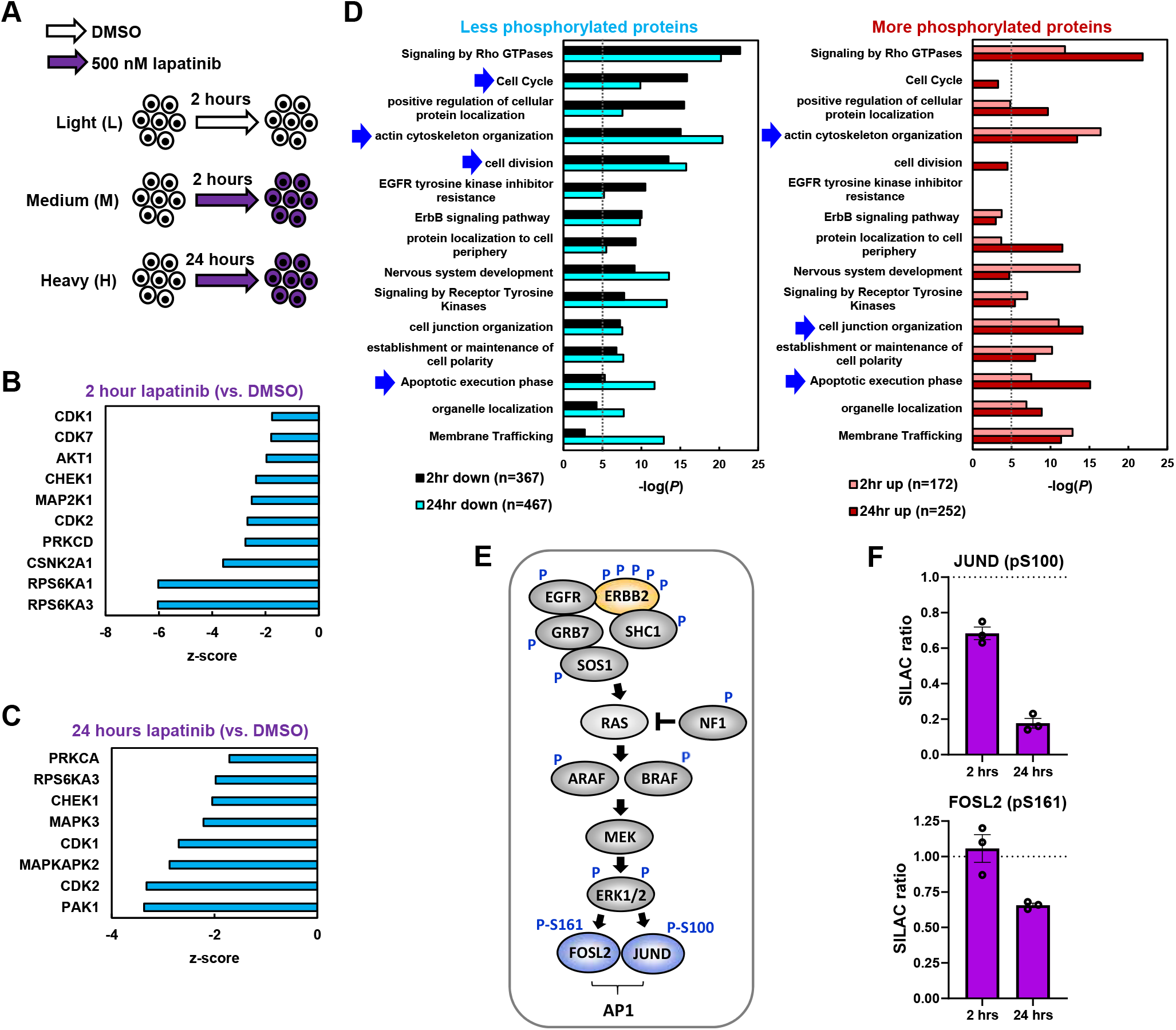
Phosphoproteomic analysis of ERBB2 signalling in OAC. (A) Schematic of mass spectrometry experimental outline. SILAC labelled OE19 cells were treated with DMSO for 2 hours, or 500 nM lapatinib for 2 and 24 hours. (B and C) Kinase set enrichment analysis predicted changes in kinase activity in OE19 cells treated with 500 nM lapatinib for (B) 2 hours or (C) 24 hours, relative to DMSO vehicle control. (D) GO analysis of genes encoding proteins that were differentially phosphorylated in OE19 cells following 2 or 24 hours lapatinib treatment relative to DMSO control. Arrows highlight terms discussed in the text. (E) Summary of phosphorylation site (P) changes after 2 and/or 24 hrs in proteins downstream from ERBB2 in the RAS-ERK pathway. (F) SILAC ratios of the peptides containing the indicated phosphorylated residues in OE19 cells treated with 500 nM lapatinib for the indicated timepoints, relative to DMSO vehicle control.

To gain insights into the impact of ERRB2-mediated phosphorylation events we analysed the local sequence context of the phosphorylated sites to determine the likely upstream kinases affected. Downregulation of the activity of multiple predicted kinases was revealed from the enriched motifs showing reduced phosphorylation, including elements of the signalling pathway through ERK, with MAP2K1 (MEK1), RPSP6KA1 (RSK1) and RPS6KA3 (RSK2) down after 2 hrs (Fig. 1B) and MAPK3 (ERK1) down at 24 hrs (Fig. 1C). Cyclin-dependent kinase activity including CDK1 and 2 were downregulated at both time points (Fig. 1B and C), consistent with the loss of cell proliferation observed following lapatinib treatment of OE19 cells (9). We also examined the molecular and cellular processes that are likely affected, by performing Gene Ontology (GO) term analysis of proteins showing decreased or increased phosphorylation following ERBB2 inhibition (Fig. 1D). Decreased phosphorylation was associated with cell cycle regulation and actin cytoskeleton organisation at both time points. Actin filament based processes were associated with increased phosphorylation as was cell junction organisation. Terms associated with apoptosis were associated with both hyper and hypophosphorylated proteins, which is consistent with the cell death observed upon lapatinib treatment of OE19 cells (9).

Subsequent analysis of the phosphorylation events in the canonical pathway downstream from ERBB2, revealed multiple changes to pathway components including phosphorylation sites on ERBB2 itself (Fig. 1E; Supplementary Fig. S1G; Supplementary Table S2). Several of the sites lost following ERBB2 inhibition are known to be important for controlling ERBB2 activity either positively (Y847; referred to as Y882 in (49)) or negatively (T671; referred to as T677 in (50)). Interestingly, several downstream transcriptional regulatory proteins show decreased phosphorylation including FOXK1/2, HMGA1, ZNF609 and the AP-1 components JUND and FOSL2 (Fig. 1E and F; Supplementary Table S2). In the case of JUND, the downregulated site (S100) is known to be important for ERK-mediated activation of its activity (51). These targets provide potential links from ERBB2 to downstream changes in gene expression.

Inhibition of ERBB2 activity is therefore associated with large changes in the cellular phosphoproteome, including multiple downstream components of the RAS-ERK pathway. Phosphorylation of several transcriptional regulators, including AP-1, is disrupted and multiple protein targets of ERBB2 signalling are involved in the key cellular processes of cell division, actin cytoskeleton organisation and apoptosis. Thus we generated a picture of the molecular events and associated cellular processes controlled by ERBB2 in OAC.

### ERBB2 regulates AP-1 activity in ERBB2-positive OAC cells

Having established the signalling pathway changes elicited by ERBB2 inhibition, we next interrogated the nuclear consequences of ERBB2 inhibition by examining its influence on the accessible chromatin landscape. Previously, we performed ATAC-seq analysis to uncover accessible chromatin regions, which revealed the regulators and processes induced following ERBB2 inhibition that are important for drug resistance (9). Here we analyse the consequences of ERBB2 inhibition in terms of downregulated regulatory activities by focussing on chromatin regions that close following ERBB2 inhibition. We identified 1481 regions opening and 824 regions closing (>2 fold linear fold change, FDR < 0.05) following 24 hrs of lapatinib treatment (9; Fig. 2A). Importantly, there was a large overlap with chromatin accessibility changes elicited by genetic depletion of *ERBB2*, which is particularly noticeable in the closing regions following more extended lapatinib treatment for 7 days (Supplementary Fig. S2A-C). These chromatin accessibility changes following ERBB2 downregulation or inhibition correlated with changes in histone H3K27ac levels, indicating that chromatin closing is associated with the loss of this activating mark and vice versa for chromatin opening (Fig. 2A; Supplementary Fig. S2D). KLF transcription factor binding motifs are enriched in both opening and closing regions (Fig. 2B and C; Supplementary Table S3A and B), consistent with the observation that we can detect KLF5 binding in both sets of regions (Fig. 2A). Regions exhibiting increased opening, showed enrichment for motifs recognised by a set of transcription factors such as HNF4A and GATA4 that are usually expressed in gastrointestinal tissues (Fig. 2B; Supplementary Table S3B). These motifs are also observed as cells acquire inhibitor resistance at later treatment time points (9). Indeed, these regions also exhibit increased HNF4A binding following lapatinib treatment (Fig. 2A). In contrast, the motif recognised by AP-1 transcription factors (TGA^C^/_G_TCA) is the dominant sequence found in regions that close following ERBB2 inhibition (~55% of regions; Fig. 2C; Supplementary Table S3A), with motifs for ETS transcription factors being the next most common occurrence (~25%; Fig. 2C; Supplementary Table S3A). HNF4A motifs are lacking, consistent with the low levels of HNF4A binding observed in the closing regions (Fig. 2A). Similar results were obtained when ERBB2 was depleted using siRNA (Supplementary Fig. S2D-F). Both AP-1 and ETS transcription factors are established transcriptional and direct post-transcriptional targets of ERK, one of the key downstream kinases activated by ERBB2 signalling (52). Importantly, the AP-1 motif was also identified as the top scoring motif in chromatin regions that closed in response to lapatinib treatment in a range of additional gastro-oesophageal adenocarcinoma cells lines and an OAC organoid which harbour ERBB2 amplifications (Fig. 2D).

**Figure 2.**
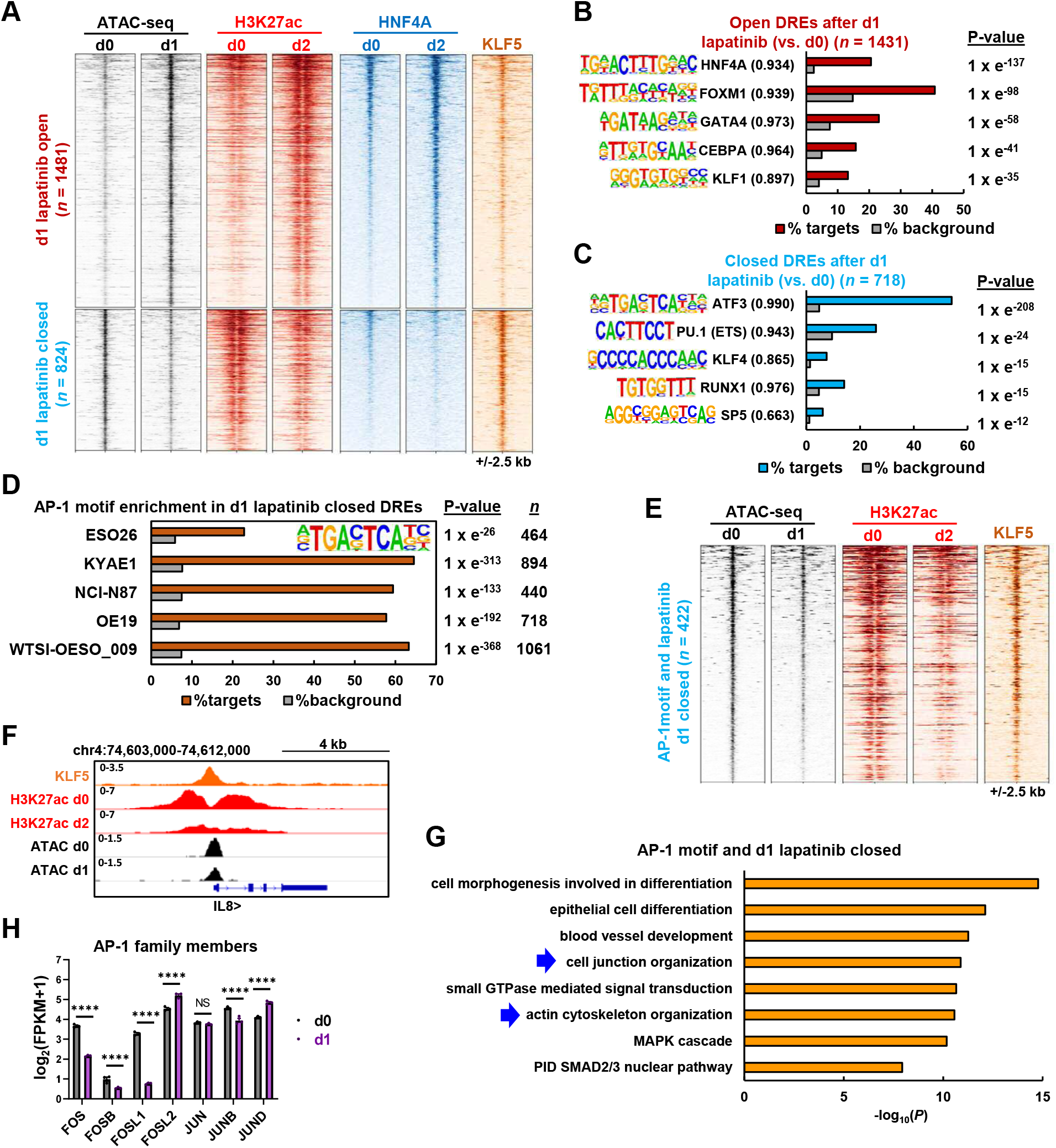
ERBB2 regulates AP-1 activity in ERBB2 positive OAC cells. (A) Heatmap of differentially open or closed ATAC-seq peaks in OE19 cells treated with 500 nM lapatinib for 1 day. ATAC-seq, HNF4A ChIP-seq and H3K27ac ChIP-seq data is shown for OE19 cells treated with lapatinib for the indicated timepoints (d0-d2). KLF5 ChIP-seq data from untreated OE19 cells is also shown. (B and C) *De novo* transcription factor motif enrichment at differentially (B) open or (C) closed distal regulatory elements (DREs) in OE19 cells treated with 500 nM lapatinib for 1 day. Motif match score to called transcription factor is shown in brackets. (D) AP-1 transcription factor motif enrichment in differentially closed DREs in ERBB2 positive gastro-oesophageal cell lines or organoid (WTSI-OESO_009) treated with 500 nM lapatinib for 1 day. (E) Heatmap of differentially closed chromatin regions containing the AP-1 motif in OE19 cells following 500 nM lapatinib treatment for 1 day (d1). ATAC-seq, H3K27ac ChIP-seq data is shown for OE19 cells treated with lapatinib for the indicated timepoints (d0-d2). Parental OE19 KLF5 ChIP-seq data is also shown. (F) Genome browser view of ATAC-seq data and ChIP-seq data (KLF5 and H3K27ac) at the indicated times of lapatinib treatment (d0, d1 or d2) highlighting the *IL8* locus. (G) GO analysis of genes annotated to 1 day lapatinib differentially closed peaks that contain the AP-1 motif. Peaks were annotated to genes by the basal extension model using GREAT, and annotated genes were then analysed using Metascape. (H) mRNA expression from RNA-seq data of OE19 cells treated with lapatinib. d0 – DMSO control, d1 – 1 day lapatinib **** *P* < 0.0001.

Given the dominance of the AP-1 motif, we focussed on the closing regions containing AP-1 motifs. These regions show decreased H3K27ac following lapatinib treatment and are generally co-bound by KLF5 (Fig. 2E), a factor we previously implicated in OAC (36). The *IL8* locus is one such example (Fig. 2F). Again, similar findings were made in AP-1 motif containing regions identified from ERBB2 depleted cells (Supplementary Fig. S2G). These regions are associated with genes enriched in GO terms related to actin cytoskeleton organisation and cell junction organisation (Fig. 2G). These terms are consistent with the processes affected at the post-translational level following ERBB2 inhibition (Fig. 1D) and highlight a potential role for AP-1 in this context. Importantly, many AP-1 family members including *FOS, FOSL1* and *JUNB* show downregulation at the RNA level following either ERBB2 inhibition by 24 hrs of lapatinib treatment (Fig. 2H) or *ERBB2* depletion (Supplementary Fig. S2H).

Collectively, these data demonstrate that pharmacological or genetic inhibition of ERBB2 signalling leads to substantive chromatin closing, and these regions are associated with AP-1 motifs, implying a loss of AP-1 activity. Indeed, reductions in AP-1 transcription factor expression following ERBB2 inhibition is consistent with this conclusion, coupled with the reductions in AP-1 component activating phosphorylation events we observe.

### AP-1 activity is elevated in ERBB2 positive OAC patients

Given the links we uncovered between ERBB2 signalling and AP-1 in our cell line models, we sought evidence for elevated AP-1 activity in OAC patients with tumours containing *ERBB2* amplifications. Our previous work based on cell line models and a limited number of patient OAC samples, used ATAC-seq to implicate AP-1 in OAC (17, 19). To more precisely study the potential role of ERBB2 in signalling through AP-1 in OAC, we compared the accessible chromatin landscape of three *ERBB2* amplified OAC tumours to Barrett’s oesophageal samples (the precursor to OAC) and nine OAC samples lacking such amplifications (Supplementary Fig. S3A). Using the three *ERBB2* amplified OAC samples, we called differentially open peaks compared to Barrett’s oesophagus (2X linear fold change, FDR < 0.05) and identified 1,253 more accessible and 679 less accessible regions (Fig. 3A; Supplementary Table S4C and D). The more accessible regions showed increased accessibility when compared to other OAC samples, suggesting an influence of *ERBB2* amplification (Fig. 3A; Supplementary Fig. S3B). Motif enrichment analysis identified the AP-1 motif (represented by FRA1) as the top scoring motif in the regions showing increased accessibility in *ERBB2* amplified OAC with ~30% of regions containing this motif (Fig. 3B; Supplementary Table S4F). Conversely, the closing regions showed an enrichment for CTCF as the top scoring motif (Fig. 3C; Supplementary Table S4F). Furthermore, similarities between motifs identified in differentially accessible regions in ERBB2-positive patients with those identified in differentially accessible regions following ERBB2 inhibition/depletion *in vitro* were uncovered (Fig. 3D). Clustering of the enriched motifs revealed the close concordance of AP-1 motif association with more accessible regions in ERBB2-positive OAC cancer samples and regions that closed in OE19 cells after ERBB2 inhibition (Fig. 3D; Supplementary Table S4G). Conversely, FOX and GATA factors showed reciprocal behaviour and were more enriched in regions more open in Barrett’s oesophagus and which opened following ERBB2 inhibition in OE19 cells (Fig. 3D; Supplementary Table S4G). GO term analysis revealed that genes associated with opening regions were enriched for a variety of biological processes related to epithelial cell biology, but also “response to EGF” and “actin cytoskeleton organisation” (Fig. 3E). The former would be expected due to the influence of amplified *ERBB2*, while the latter process is in line with the changes observed in the phosphoproteome (Fig. 1D).

**Figure 3.**
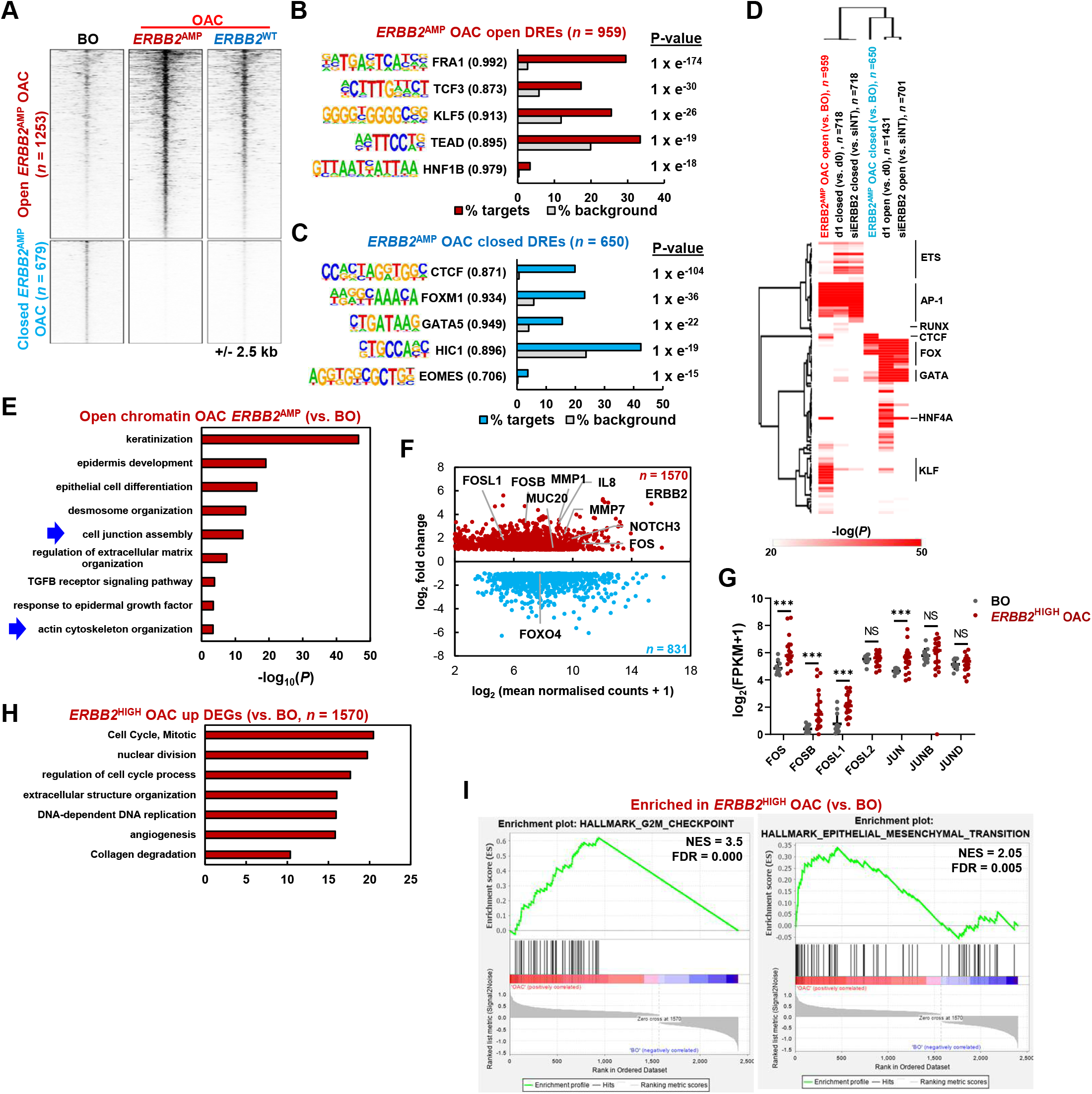
AP-1 expression increases during the transition from Barrett’s oesophagus to ERBB2 positive oesophageal adenocarcinoma. Heatmap showing ATAC-seq signal at differentially open or closed (FDR < 0.05) chromatin regions in OAC tumours harbouring *ERBB2* amplifications (*ERBB2*^AMP^ OAC) relative to Barrett’s oesophagus (BO) tissue. Data for OAC tumours without *ERBB2* amplifications (*ERBB2*^WT^ OAC) is also shown. (B and C) *De novo* transcription factor motif enrichment at differentially (B) open or (C) closed distal regulatory elements. Motif match score to called transcription factor is shown in brackets. (D) Heatmap of Homer ‘known’ transcription factor motif enrichment (log_10_*P* > 20 in one condition) in chromatin regions closing or opening in OE19 cells with lapatinib (d1) or siERBB2 treatment or in patients harbouring ERBB2 amplifications (ERBB2+) compared to Barrett’s oesophagus (BO) patients. (E) GO analysis (GREAT) of genes annotated (basal extension model) to differentially open chromatin regions in *ERBB2*^AMP^ OAC relative to BO. (F) MA plot of differentially expressed genes (DEGs) in *ERBB2*^HIGH^ OAC patient samples relative to Barrett’s oesophagus tissue. Red – genes up-regulated in *ERBB2*^HIGH^ OAC; blue – genes down-regulated in *ERBB2*^HIGH^ OAC. DEGs were defined by FDR < 0.05, 2X fold change and FPKM > 1. (G) mRNA expression of AP-1 family members in Barrett’s oesophagus (BO) and *ERBB2*^HIGH^ OAC patient tissue. (H) GO analysis (Metascape) of genes up-regulated in *ERBB2*^HIGH^ OAC relative to BO tissue. (I) Enrichment plots from GSEA on differentially expressed up- and down-regulated genes in *ERBB2*^HIGH^ OAC relative to BO. NES - normalised enrichment score.

To further examine the molecular changes in OAC samples containing high levels of *ERBB2* expression (defined as >2SD above the median level), we compared their RNAseq profiles with those found in Barrett’s oesophagus patients. PCA analysis clustered each type of sample together (Supplementary Fig. S3C) and we uncovered 1,570 upregulated and 831 downregulated genes (Fig. 3F; Supplementary Table S4A and B). Among the upregulated genes we identified the AP-1 family genes *FOS, FOSB* and *FOSL1* (Fig. 3F and G). GO term analysis revealed an increase in cell cycle-associated genes as might be expected for cancer cells (Fig. 3H) and decreases in genes associated with various metabolic processes (Supplementary Fig. S3D). However, we also identified “collagen degradation” as an enriched term associated with up regulated genes, and further analysis of our ATAC-seq data revealed prominent peaks in the intragenic region associated with the *MMP7* and *MMP20* genes encoding matrix degrading enzymes (Supplementary Fig. S3E). RNAseq analysis revealed upregulation of genes encoding several MMPs in these OAC samples including *MMP7* (Fig. 3F; Supplementary Fig. S3F). Finally, we looked for Gene Set enrichments in our data and again identified various cell cycle terms associated with the upregulated genes, but also “epithelial to mesenchymal transition (EMT)” (Fig. 3I; Supplementary Fig. S3G). Metabolic processes were again associated with downregulated genes (Supplementary Fig. S3G).

Collectively, these data therefore support a potential functional role for AP-1 in OAC patient samples exhibiting high level *ERBB2* expression, both from upregulation of its constituent subunits and the appearance of its binding motif in opening regions of chromatin. Furthermore, genes associated with biological processes focussed around cell cycle, actin cytoskeletal rearrangement, ECM degradation and EMT are prominent in these OAC samples, suggesting a potential link between these molecular and gene expression events.

### AP-1 regulates processes associated with EMT in ERBB2 positive OAC cells

Our ATAC-seq and RNA-seq data from OAC patients and cell lines and accompanying phosphoproteomic data, all point to a potentially important connection between AP-1 and the OAC cell phenotype driven by ERBB2. To determine how AP-1 affects OAC function, we inhibited AP-1 function by inducibly overexpressing a dominant-negative form of FOS (dnFOS; Fig. 4A)(18) in OE19 cells. dnFOS expression gave the expected downregulation of the AP-1 target gene *MMP1* (Supplementary Fig. S4A). However, unexpectedly, given previous links between AP-1 and cell cycle control (15; reviewed in 14), there was little effect on cell growth even 6 days after dnFOS induction (Fig. 4B). Similarly dnFOS barely affected the growth of two other ERBB2 positive OAC lines (ESO26 and KYAE1; Supplementary Fig. S4B). Indeed, there was little effect on the expression of a panel of genes related to proliferation and cell cycle control although the control AP-1 target gene *MMP1* was downregulated (Supplementary Fig. S4C). To provide further insights into the function of AP-1 in OAC, we therefore performed RNAseq analysis on OE19 (28) and KYAE1 cells after expression of dnFOS for 2 days. RNAseq replicates showed good correlation (Supplementary Fig. S4D and E). We next identified genes that were differentially expressed in the presence of dnFOS and identified 168 upregulated and 340 downregulated genes in OE19 cells and 395 upregulated and 448 downregulated genes in KYAE1 cells (Fig. 4C; Supplementary Table S5). Importantly there was a significant large overlap in down regulated targets with 171 genes (>50%) showing downregulation in both cell lines (Fig. 4D) compared to only 49 genes (<30%) showing consistent upregulation (Supplementary Fig. S4F). Ingenuity Pathway Analysis of RNA-seq data revealed the expected inhibition of activity of the AP-1 family members FOS, FOSL1 and JUN following dnFOS expression in both OE19 and KYAE1 cell lines (Figure 4E). The dnFOS-mediated downregulated genes were associated with a number of GO terms related to EMT and cell migration such as “response to wounding”, “regulation of cell adhesion” and “actin cytoskeleton organisation” (Fig. 4F). Example genes related to these GO terms include *MMP7, ITGB4, IL8* and *SMAD3* (Supplementary Fig. S4G). In contrast, no changes were observed in *CCND* gene expression, consistent with a lack of a connection to cell cycle and proliferation categories (Supplementary Fig. S4H). No GO terms were identified at the same levels of significance for the upregulated genes (Supplementary Fig. S4I). Gene set enrichment analysis (GSEA) revealed EMT as a highly scoring category for genes downregulated in both cell lines (Fig. 4G; Supplementary Fig. S4J).

**Figure 4.**
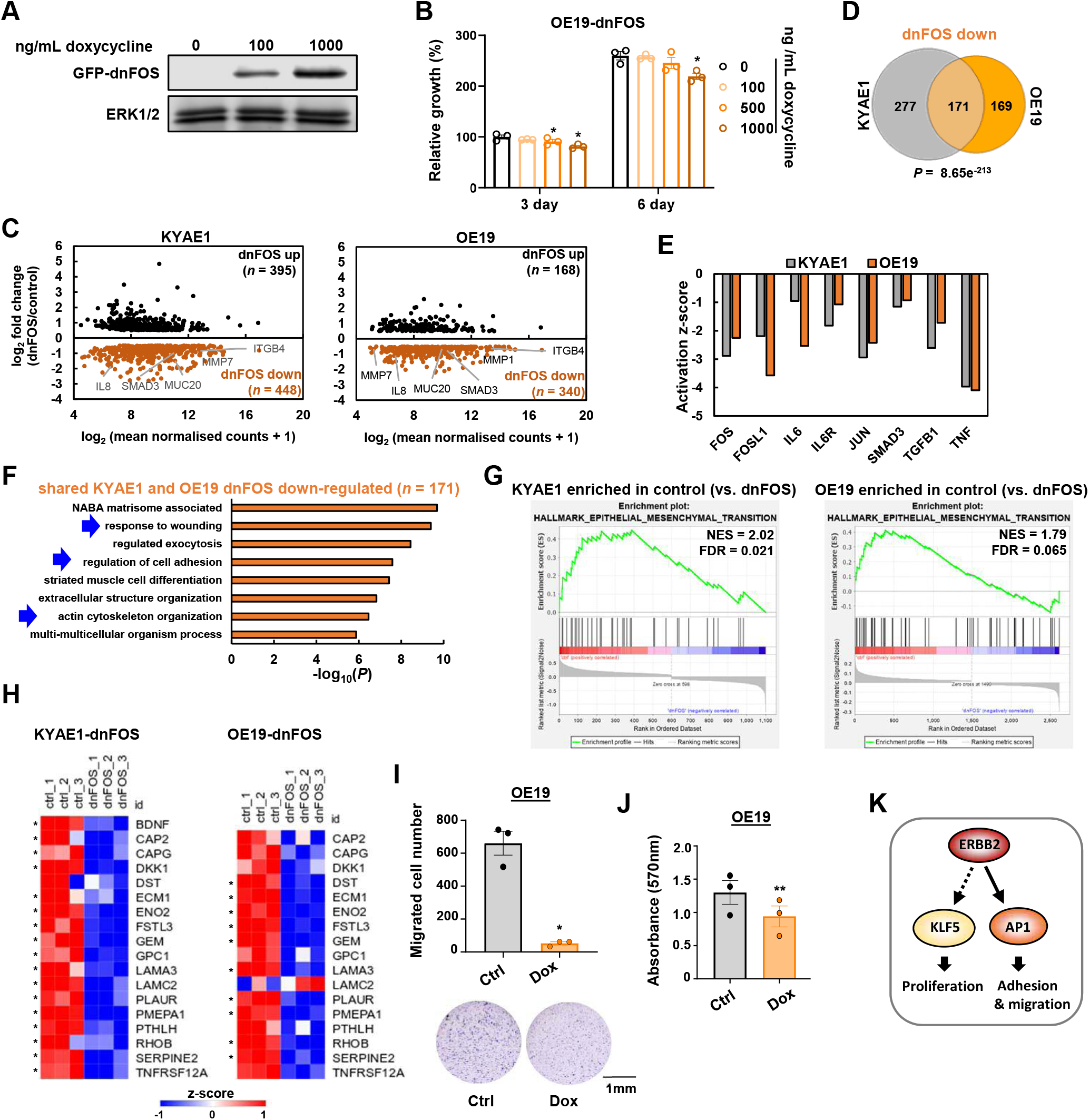
AP-1 regulates processes associated with epithelial-mesenchymal transition in ERBB2 positive OAC cells. (A)Western blot of GFP-dnFOS protein levels in OE19-dnFOS cells. GFP-dnFOS expression was induced by treatment with the indicated doses of doxycycline for 48 hours. (B) Crystal violet growth assay of OE19-dnFOS cells following dnFOS induction. * *P* < 0.05, paired T-test (*n* = 3). (C) MA plot showing DEGs (0.5X log_2_ fold change, FDR < 0.05, FPKM > 1) in KYAE1-dnFOS and OE19-dnFOS cells following GFP-dnFOS induction. Cells were untreated (control) or dnFOS expression was induced by 1000 ng/mL doxycycline treatment for 48 hours (dnFOS). (D) Overlap of genes down-regulated (log_2_ fold change > 0.5, FDR < 0.05, FPKM > 1) by dnFOS induction in KYAE1-dnFOS and OE19-dnFOS cells (i.e. shared AP-1 target genes). (E) Ingenuity Pathway Analysis of proteins whose activity is predicted to be inhibited in KYAE1-dnFOS and OE19-dnFOS cells following dnFOS induction. (F) GO analysis (Metascape) of shared AP-1 target genes. (G) Enrichment plot of epithelial-mesenchymal transition hallmark from GSEA on differentially expressed up- and down-regulated genes in OE19 cells following dnFOS induction. NES - normalised enrichment score. (H) Z-scored mRNA expression of partial EMT (pEMT) genes (34) down-regulated in either KYAE1-dnFOS or OE19-dnFOS cells following dnFOS induction. * Denotes statistically significant changes (0.5 log_2_-fold change, FDR < 0.05). Note that *BDNF* expression was only detected in KYAE1 cells. (I) Cell migration assay of OE19-dnFOS cells treated with DMSO (ctrl) or doxycycline (Dox). ** *P* < 0.01, paired T-test (*n* = 3). (J) Adhesion assay of OE19-dnFOS cells treated with DMSO (ctrl) or doxycycline (Dox). ** *P* < 0.01, paired T-test (*n* = 3). (K) Model for ERBB2 signalling through AP-1 and KLF5 to promote distinct biological processes. Dotted arrow indicates that direct molecular connections to KLF5 have not been shown (35).

Recently, a partial EMT-like state (pEMT) was identified as intrinsic to epithelial-derived cancer cells rather than the stromal compartment (33). We therefore asked whether expression of any of the 79 genes in the OAC pEMT signature are affected by AP-1 inhibition. 17 of the pEMT genes were significantly down regulated in either KYAE1 or OE19-dnFOS cells, with the majority of these showing downregulation in both cell lines (Fig. 4H) with only one gene going in the opposite direction in both cases. This finding is substantiated by GSEA using a custom dataset created from the 79 pEMT genes (Supplementary Fig. S4K). From the 17 pEMT genes responsive to AP-1 inhibition, 8 of these show upregulation in ERBB2^HIGH^ OAC patient samples, while none showed lower expression (Supplementary Fig. S4L). This is consistent with notion that this AP-1-regulated pEMT signature is active in ERBB2-driven OAC patients. To investigate further links to ERBB2 signalling, we also tested overlaps between the AP-1 target genes and those regulated by ERBB2 inhibition by genetic depletion or pharmacological inhibition. In both cases a significant overlap was seen between downregulated genes when a stringent 2 fold cut off was applied (Supplementary Fig. S4M) or when we considered the directionality of response (Supplementary Fig. S4N). These results are therefore consistent with a link between ERBB2 signalling and AP-1 activity.

Finally, we studied the functional consequences of AP-1 inhibition based on the predictions from enriched GO terms and gene sets. First, we tested cell migration following dnFOS treatment of OE19 and KYAE1 cells and found a significant decrease in both cell lines (Fig. 4I; Supplementary Fig. S4O). We also assessed cell adhesion and found it to be severely disrupted in both lines (Fig. 4J; Supplementary Fig. S4P).

Collectively these data therefore support a role for AP-1 in mediating the outputs from oncogenic ERBB2 signalling through affecting gene expression and the cellular functions associated with EMT-like processes.

## Discussion

Signalling through the RAS-ERK pathway is a hallmark of OAC and genetic disruptions to this pathway are observed in a large proportion of patients, indicating that is likely a pivotal event in the tumourigenic process (4,5). The most frequent aberrations observed are amplifications of the gene encoding the upstream RTK, ERBB2 and here we focussed on patients and cell lines harbouring this amplification event to gain mechanistic insights into the pathways through which ERBB2 controls OAC cell behaviour. Through combined analysis of phosphoproteomic, open chromatin and transcriptomic datasets we converged on a model whereby ERBB2 functions through activating the transcription factor AP-1 and its downstream target gene network to promote mesenchymal-like behaviour through changes to cell adhesion and migration (Fig. 4K).

Functionally, we link AP-1 activity in OAC cells to control of EMT-related processes such as actin cytoskeletal structures, cell migration and cell adhesion. This is consistent with the functions of AP-1 in other cancer subtypes such as the recently uncovered link between AP-1 and mesenchymal transformation in glioblastomas (53) and more broadly links between AP-1 and mesenchymal-like behaviour across multiple cancer types (54). More generally, ERBB2 is also involved in these processes as exemplified from our phosphoproteomics and open chromatin profiling experiments following genetic or chemical inhibition of ERBB2 in OAC cells. Thus, the molecular links we make are supported by the functional similarities we uncover. Our data are consistent with earlier studies that linked AP-1 to RTK signalling in other cancer types where similar biological processes are affected (55). However, we were unable to uncover any links to *CCND* expression or more generally cell cycle and proliferation control, suggesting that ERBB2 controls this aspect of cell behaviour through alternative regulatory proteins. The transcription factor KLF5 is one such candidate, which we recently linked to cell cycle control in OAC (Fig. 4K; 36).

Our phosphoproteomic studies show that a multitude of phosphorylation changes are initiated shortly after ERBB2 inhibition and that this rewiring is maintained or further modified after 24 hrs. While we have focussed on the RAS-ERK pathway and the functional downstream consequences from this, many other kinase activities are altered. Central among these are the cyclin-dependent kinases (CDKs) whose activities are downregulated and likely contributes to the proliferation defects caused by ERBB2 inhibition. However, AKT1 and PRKCD (PKCδ) are among the immediately affected kinases and PAK1 features prominently after 24 hrs (Fig. 1B and C). The latter kinase is of interest as it lies downstream of Cdc42/Rac signalling and transmits signals to the actin cytoskeleton reorganisation and subsequent cell motility. It is also defined as a “helper gene” and is amplified in ~4% of OAC cases (56). Thus, ERBB2 may impact both directly through modifying signalling cascade activity and indirectly through altering the cellular transcriptome and proteome, leading to rewiring of the signalling networks in the cell and subsequent phenotypic effects. Further work is required to understand the full impact of ERBB2 regulated pathways on the cellular behaviour of OAC cells.

While we have shown that AP-1 is the dominant transcriptional regulator downstream of ERBB2 signalling, it is likely that additional proteins are involved controlling ERBB2-mediated gene expression programmes. In addition to AP-1, our data implicate several other potential transcriptional regulators in ERBB2 functionality in OAC. For example, the changes in phosphorylation status of FOXK1/2, HMGA1, and ZNF609 suggest likely downstream gene regulatory consequences. Furthermore, open chromatin profiling reveals loss of accessibility in sites enriched for the binding of ETS, KLF and RUNX transcription factors following ERBB2 inhibition. Members of the ETS family (17, 57, 58) and KLF5 (36, 58) have previously been implicated as important regulators of OAC gene expression programmes. *RUNX2* has been identified as a helper gene and is amplified at a low level in OAC (~2%; 56). Additional transcriptional regulators may have been missed by our analyses either due to the sensitivity of the phosphoproteomic analysis or for proteins whose binding site accessibility does not greatly change (eg by transcription factor exchange) and/or have relatively few target loci. Again further work is required to gain a comprehensive view of the transcriptional regulatory proteins controlled by oncogenic ERBB2 signalling.

In summary, we demonstrate that oncogenic ERBB2 signalling controls cell behaviour through modifying the activity of the AP-1 transcription factor. AP-1 itself drives processes associated with cell migration and invasion which might ultimately contribute to the metastatic spread of the tumours. Importantly, our results have broader significance across a wider range of OAC patients, given the frequent activation of other RTKs and downstream pathway components (4, 5), which likely also impact on AP-1 activity. ERBB2 inhibition has been shown to have limited clinical efficacy with only a few months gained in overall survival time (6, 59). Instead, targeting components of the downstream AP-1 regulatory network might represent an alternative more effective therapeutic route through which to combat ERBB2-driven OAC.

## Acknowledgements

We thank Guanhua Yan for excellent technical assistance, staff in the Bioinformatics, Genomic Technologies and Bioimaging core facilities. We also thank Angeliki Malliri, and Shen-Hsi Yang for critical appraisal of the manuscript. We thank Joseph Parsons and Dr David Knight (Bio-MS Facility) for help with the phosphoproteomics experiments. This work was funded by grants to ADS from the Wellcome Trust (103857/Z/14/Z, 102171/Z/13/Z and 102171/B/13/Z). Research in CF lab is supported by the Wellcome Trust (107636/Z/15/Z and 107636/Z/15/A).

## Author contributions

S.O. and I.A. performed the experiments and data analysis in this study; P.F. prepared samples for proteomics and phosphoproteomics analysis. C.F. led the proteomic studies, performed data analysis and supervised P.F. and S.O.; A.D.S. led the other experimental parts of the program. All authors contributed to manuscript preparation and/or critically appraised manuscript drafts.

---

**Figure S1.**
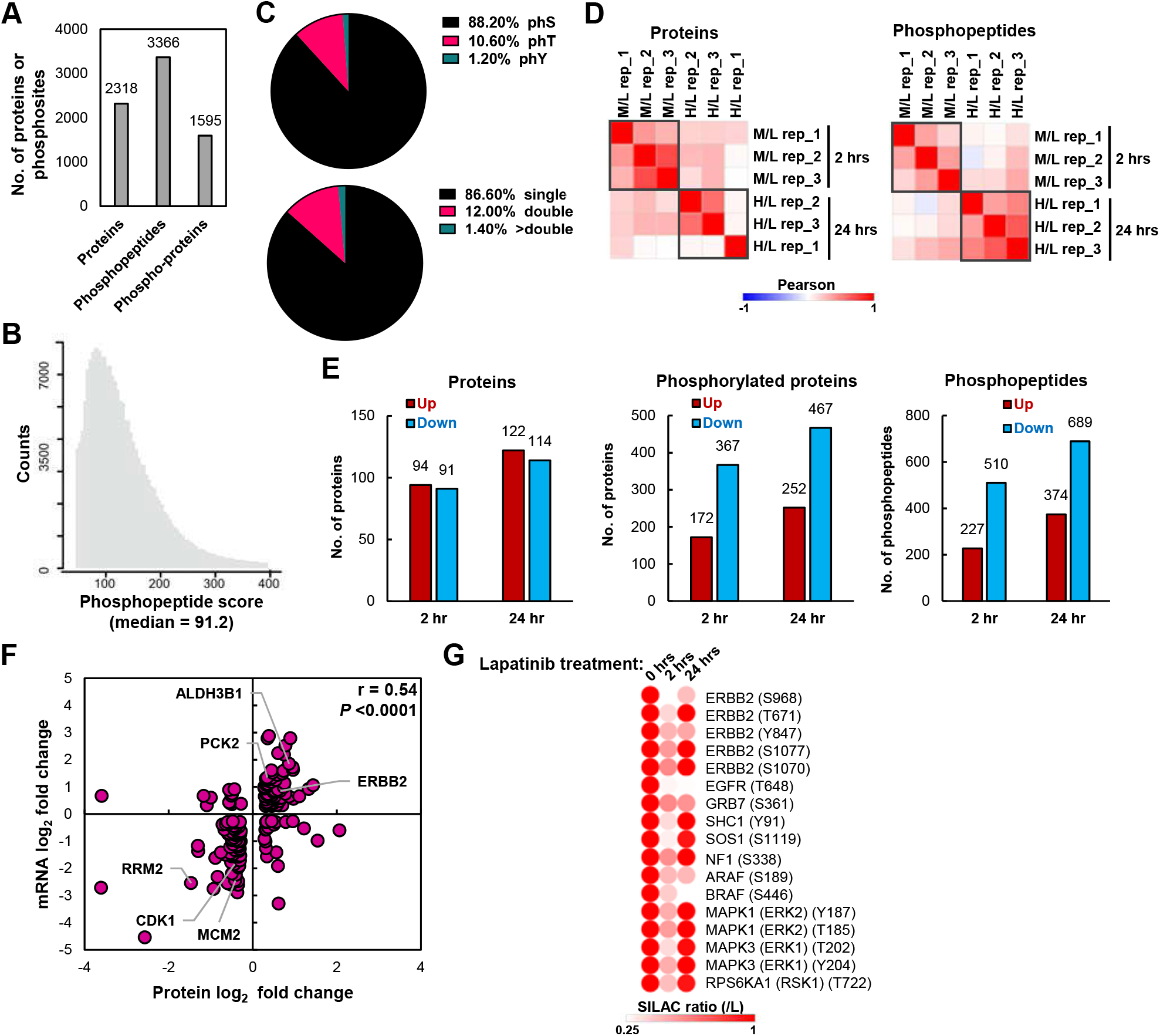
Phosphoproteomic analysis of ERBB2 signalling in OAC cells. (A) The number of proteins, phosphopeptides and phosphorylated proteins detected in all 3 biological replicates across all conditions. (B) Phosphorylated peptide score. (C) Distribution of phosphorylated sites; phS – phosphoserine, phT – phosphothreonine, phY – phosphotyrosine (top) and of phosphopeptides with 1, 2 or more phosphorylated sites (bottom). (D) Pearson correlation of proteome replicates (PRO) and phosphoproteome replicates (STY) of OE19 cells treated with 500 nM lapatinib for 2 hours or 24 hours. SILAC ratios are expressed as lapatinib treated sample relative to DMSO control. H – heavy, 24 hours lapatinib; M – medium, 2 hours lapatinib; L – light, 2 hours DMSO. (E) The number of up- or down-regulated: proteins or phosphorylated proteins or phosphopeptides after 2 or 24 hours lapatinib treatment relative to DMSO control. Up- or down-regulated proteins/phosphorylated proteins/phosphopeptides were defined based upon all 3 biological replicates having a 1.2X linear fold change in the same direction. (F) Correlation of changes in the proteome and transcriptome after 24 hours 500 nM lapatinib treatment relative to DMSO control. Only genes in which the protein and mRNA fold change was greater than 1.2 fold are shown. Pearson correlation (r) is shown. (G) Heatmap of SILAC ratios for peptides containing the indicated phosphorylation sites before (0 hr) and after (2 and 24 hr) lapatinib treatment (relative to 0 hrs).

**Figure S2.**
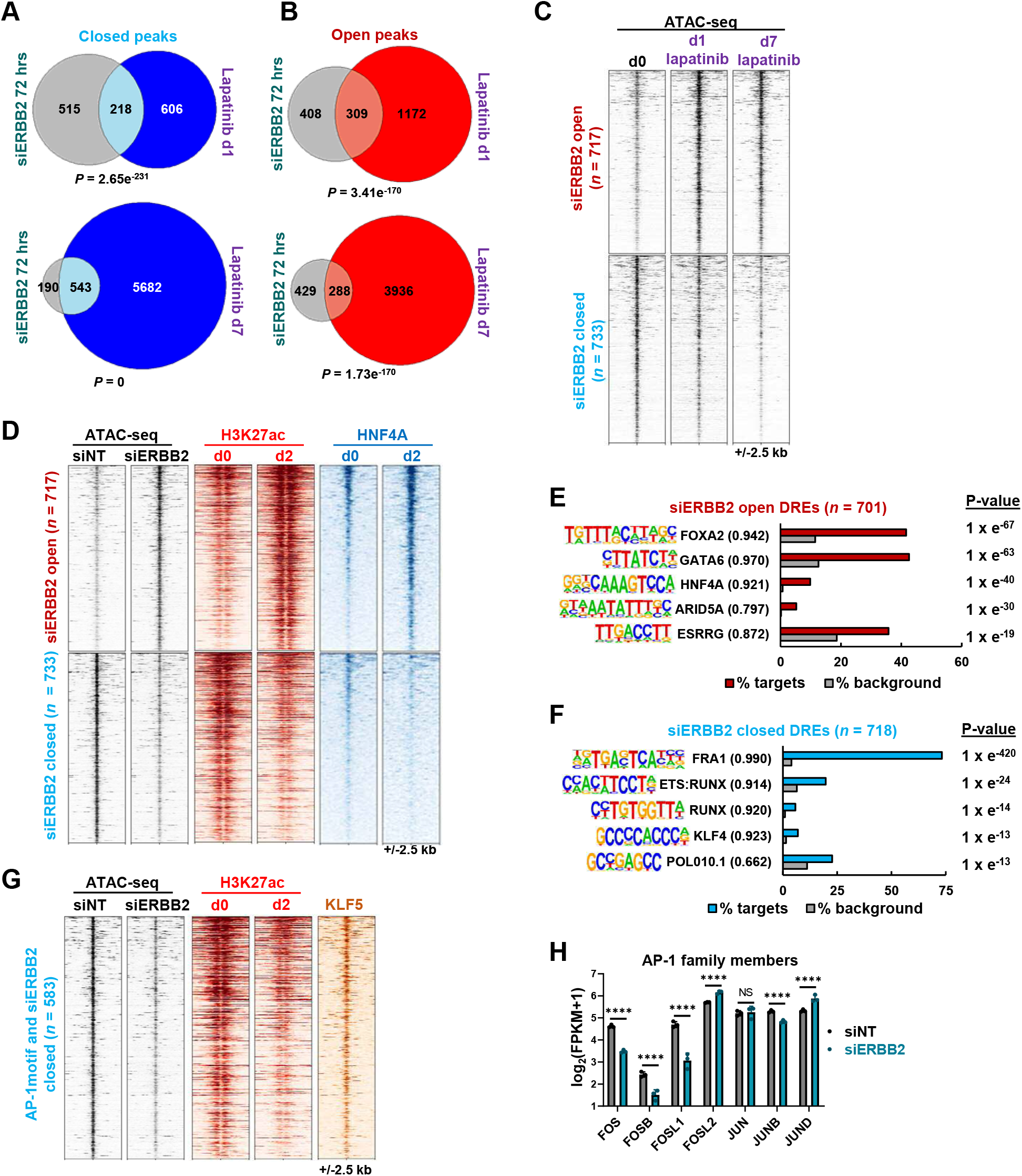
ERBB2 regulates AP-1 activity in ERBB2 positive OAC cells. (A and B) Overlap of differentially accessible peaks in OE19 cells after ERBB2 knockdown (siERBB2) and 1 (d1) or 7 (d7) day 500 nM lapatinib treatment. (A) Overlap of closed peaks. (B) Overlap of open peaks. (C) Heatmap showing ATAC-seq and signal in OE19 cells treated with lapatinib (d0-d7) at differentially open or closed ATAC-seq peaks in OE19 cells following ERBB2 knockdown (siERBB2). (D) Heatmap of differentially open or closed ATAC-seq peaks in OE19 cells following ERBB2 knockdown (siERBB2). ATAC-seq is shown for OE19 cells treated with siERBB2. HNF4A ChIP-seq and H3K27ac ChIP-seq data is shown for OE19 cells treated with lapatinib for the indicated timepoints (d0 or d2). (E and F) *De novo* transcription factor motif enrichment at differentially (E) open or (F) closed distal regulatory elements in OE19 cells following *ERBB2* knockdown. Motif match score to called transcription factor is shown in brackets. (G) Heatmap of differentially closed chromatin regions containing the AP-1 motif in OE19 cells following *ERBB2* knockdown. ATAC-seq is shown for OE19 cells treated with siERBB2. H3K27ac ChIP-seq data is shown for OE19 cells treated with lapatinib for the indicated timepoints (d0 or d2). Parental KLF5 ChIP-seq data is also shown. (H) mRNA expression of the indicated AP-1 subunits from RNA-seq of OE19 cells treated with siERBB2.

**Figure S3.**
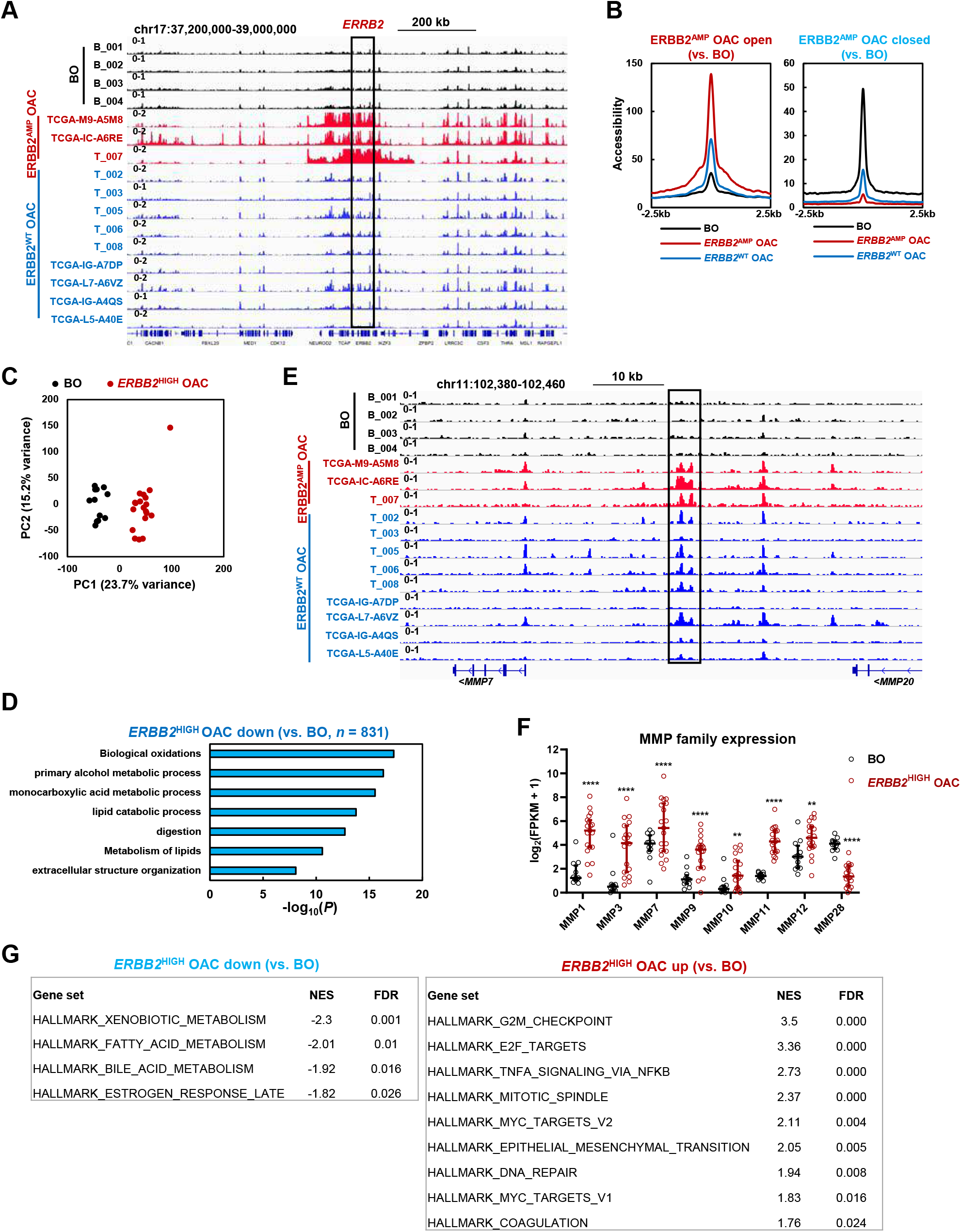
AP-1 expression increases during the transition from Barrett’s oesophagus to ERBB2 positive oesophageal adenocarcinoma. (A) Genome browser view of ATAC-seq data highlighting the *ERBB2* locus in Barrett’s oesophagus (BO) and OAC patient tissue samples. (B) Average tag density plot of chromatin accessibility measured by ATAC-seq in the indicated patient sample groups for differentially opening (left) and closing (right) regions in *ERBB2*^AMP^ OAC samples relative to BO samples (2X linear fold change, FDR < 0.05). (C) PCA of RNA-seq data from BO and *ERBB2*^HIGH^ OAC patient tissue samples. (D) GO analysis (Metascape) of genes down-regulated in *ERBB2*^HIGH^ OAC relative to BO tissue. (E) Genome browser view showing ATAC-seq signal at *MMP7* and *MMP20* loci. BO, *ERBB2*^AMP^ OAC and *ERBB2*^WT^ OAC patient tissue samples are shown. (F) mRNA expression of the *MMP* encoding genes differentially expressed between Barrett’s oesophagus (BO) and *ERBB2*^HIGH^ OAC patient tissue; ** *P* < 0.01, **** *P* < 0.0001. All are up-regulated except for *MMP28*. (G) GSEA results of differentially expressed genes in *ERBB2*^HIGH^ OAC tissue relative to BO tissue. NES – normalised enrichment score.

**Figure S4.**
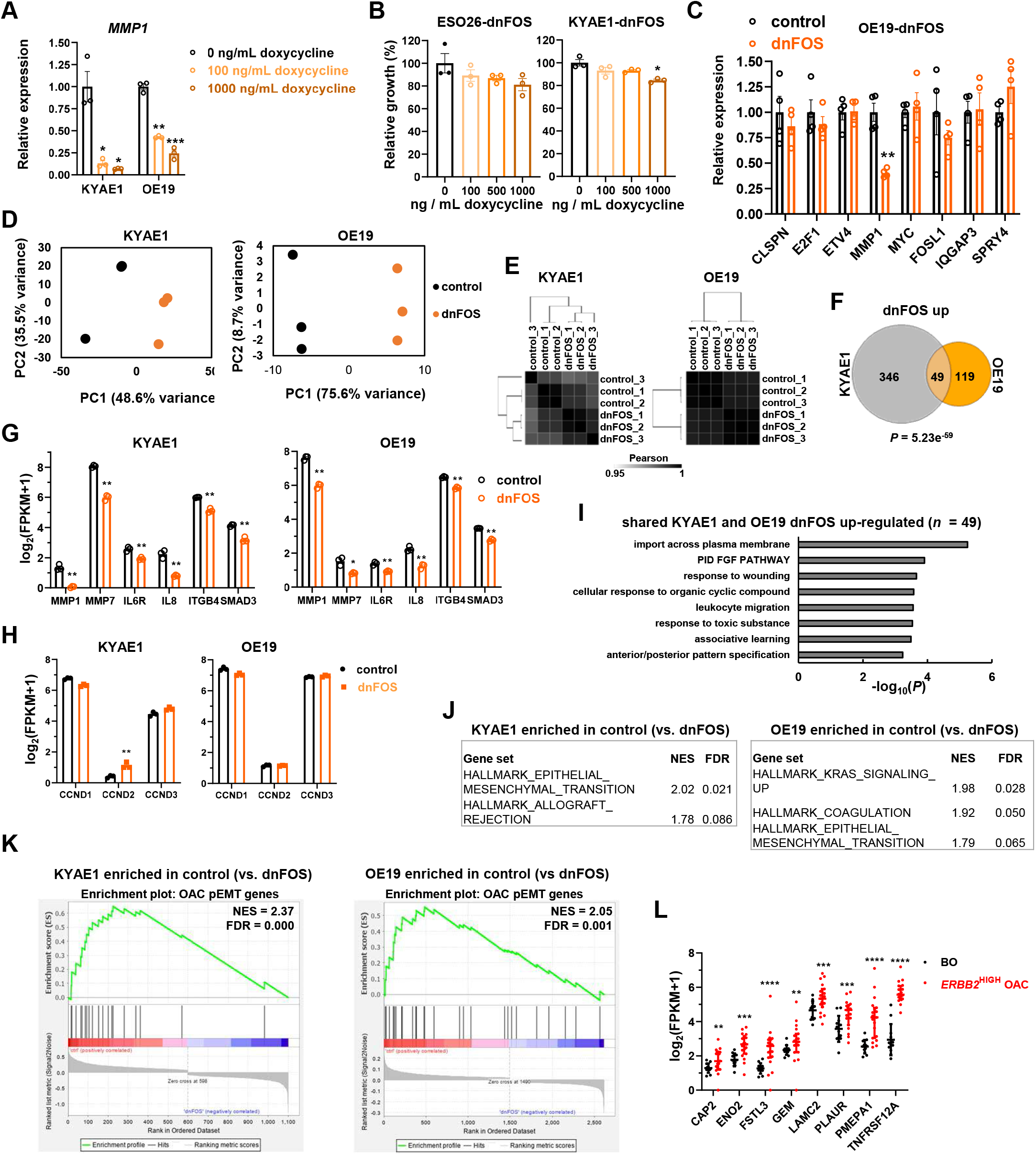
AP-1 regulates processes associated with epithelial-mesenchymal transition in ERBB2 positive OAC cells. Continued overleaf….

**Figure S4.**
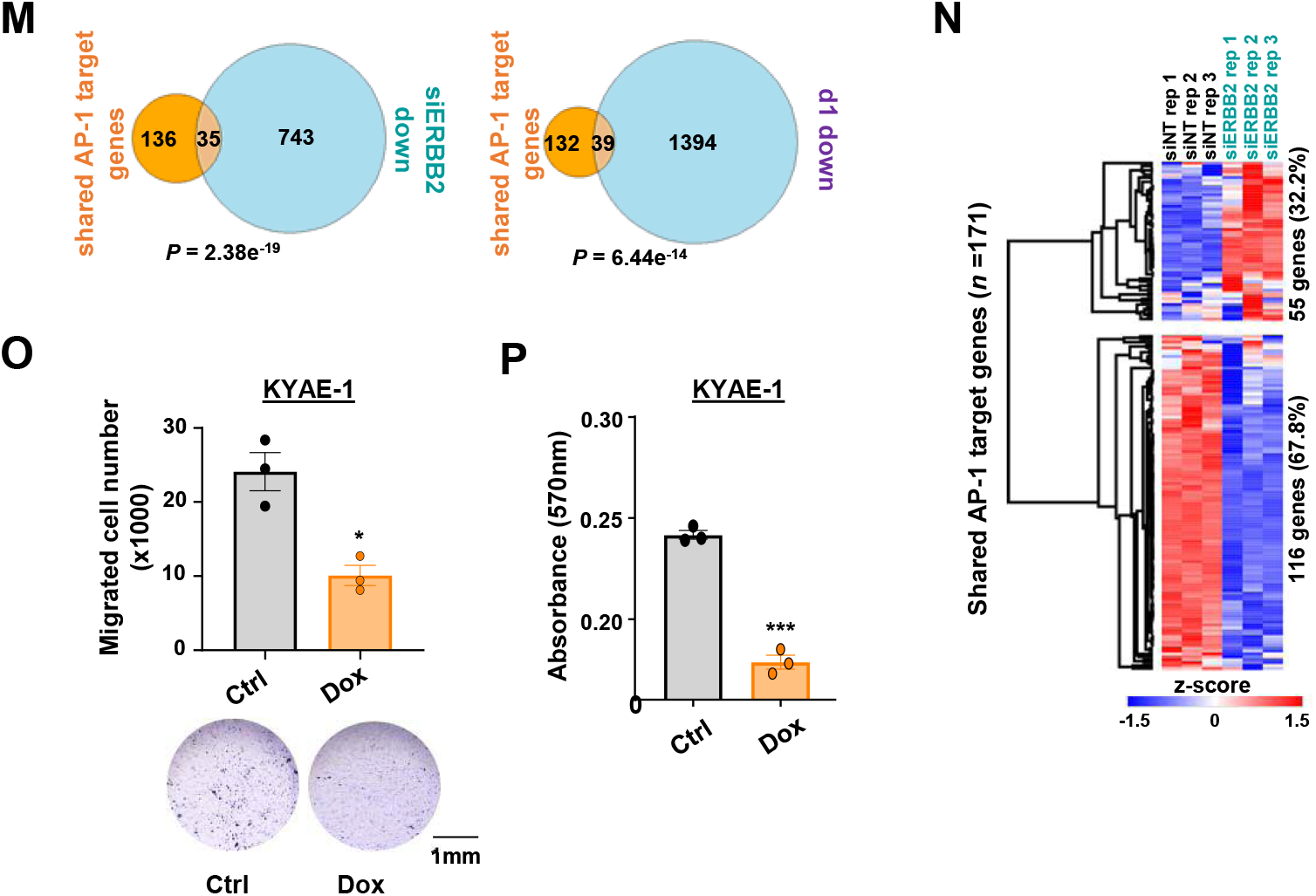
Figure S4. AP-1 regulates processes associated with epithelial-mesenchymal transition in ERBB2 positive OAC cells. (A) RT-qPCR analysis of *MMP1* expression in KYAE1-dnFOS and OE19-dnFOS cells. GFP-dnFOS expression was induced by treatment with the indicated doses of doxycycline for 48 hours. * *P* < 0.05, ** *P* < 0.01, *** *P* < 0.001; paired T-test, *n* = 3. (B) Crystal violet growth assay of ESO26-dnFOS and KYAE1-dnFOS cells following dnFOS induction. * *P* < 0.05; paired T-test, *n* = 3. (C) RT-qPCR analysis in OE19-dnFOS cells of dnFOS down-regulated genes identified in OE33 cells (Britton et al., 2017). Control – untreated cells; dnFOS – 1000 ng / mL doxycycline for 48 hours. ** *P <* 0.01; paired T-test, *n* = 4. (D) PCA and (E) Pearson correlation analysis of KYAE1-dnFOS and OE19-dnFOS RNA-seq data (*n* = 3). Control – untreated cells; dnFOS – 1000 ng / mL doxycycline for 48 hours. (F) Overlap of genes up-regulated (0.5 log_2_ fold change, FDR < 0.05, FPKM > 1) by dnFOS induction in KYAE1-dnFOS and OE19-dnFOS cells. (G and H) mRNA expression of the indicated genes in KYAE1-dnFOS and OE19-dnFOS cells from RNAseq analysis (*n* = 3). (I) GO analysis (Metascape) of genes up-regulated by dnFOS in both KYAE1 and OE19 cells. (J) GSEA results of differentially expressed genes in KYAE1 and OE19 control cells relative to dnFOS induced cells. NES – normalised enrichment score. (K) Enrichment plot from GSEA of pEMT genes (Tyler and Tirosh, 2021) on differentially expressed up- and down-regulated genes (FDR < 0.05) in KYAE1 (left) or OE19 (right) cells following dnFOS induction. NES – normalised enrichment score. (L) Patient tissue mRNA expression of pEMT genes (Tyler and Tirosh, 2021) that are down-regulated following dnFOS induction in either KYAE1 or OE19 cells. Genes up-regulated in ERBB2^HIGH^ OAC relative to BO (FDR < 0.05) are shown. (M) Overlap of shared AP-1 target genes (genes down-regulated by dnFOS in both KYAE1 and OE19 cells) and genes down-regulated (2X linear fold change, FDR < 0.05, FPKM > 1) by siERBB2 (left) or 1 day lapatinib (right) in OE19 cells. Statistical significance was determined using a hypergeometric test. (N) Heatmap showing the z-scored expression of shared AP-1 target genes in OE19 cells treated with siERBB2. The number and percentage of up- or down-regulated genes is indicated. (O) Cell migration assay of KYAE1-dnFOS cells treated with DMSO (ctrl) or doxycycline (Dox). ** *P* < 0.05, paired T-test (*n* = 3). (P) Adhesion assay of KYAE1-dnFOS cells treated with DMSO (ctrl) or doxycycline (Dox). *** *P* < 0.001, paired T-test (*n* = 3).

